# Topoisomerase IIα C-terminal Domain Mutations and Catalytic Function

**DOI:** 10.1101/2023.07.29.551120

**Authors:** Jessica R. Musselman, Daniel C. England, Lauren A. Fielding, Clay T. Durham, Emma Baxter, Xiaohua Jiang, Edward C. Lisic, Joseph E. Deweese

## Abstract

Topoisomerase IIα is a nuclear enzyme needed for dealing with topological entanglements in the DNA arising from replication and transcription. The N-terminal region and core of the protein are utilized in the catalytic cycle of the enzyme, which generates a transient double-stranded break in one segment of DNA and passes another segment through the break. The C-terminal domain is a large, intrinsically disordered region that appears to be involved in regulating the function of the enzyme both in terms of substrate selection and the level of activity of the enzyme. In a previous study, we explored eleven targeted mutations to the C-terminal domain. This present study explores six of these mutants to determine whether there are any defects in closure of the N-terminal clamp and whether an experimental compound known as a Cu(II)-thiosemicarbazone affects DNA cleavage with the mutants. Based upon our results, the mutants are able to close the N-terminal clamp, but some of the mutants that displayed the least clamp closing activity also had the lowest catalytic activity. Further, Cu-APY-ETSC did impact the ability of the enzymes to cleave DNA to similar levels as seen with the WT enzyme. These results lay the groundwork for additional analyses of the C-terminal domain and indicate the C-terminal domain regions tested did not influence the action of Cu-APY-ETSC except at the level of coordination between the two active sites.

## 1. Introduction

Topoisomerases are a family of enzymes that alter DNA topology and are found in all domains of life [1]. There are two broad families of topoisomerases: Type I and Type II. While these families differ functionally and structurally, they both participate in altering the topology of DNA using either a single-stranded (Type I) or double-stranded (Type II) DNA break [1]. Topoisomerases generate transient, reversible strand breaks to alter DNA topology during cellular activities such as DNA replication and transcription. Topoisomerases are essential cellular enzymes for all known forms of life and disruption of topoisomerase function can be lethal at the cellular level [2-4]. As a result, topoisomerases are the targets of anticancer and antibacterial drugs [2-4].

Topoisomerase II is a Type II topoisomerase that is a critical anticancer drug target in humans. Human topoisomerase II has two isoforms: α and β. Human topoisomerase IIα (TOP2A) is involved in replication and chromosome segregation, while human topoisomerase IIβ (TOP2B) is involved in chromatin regulation and transcription [1,3,4]. Based on these different roles, TOP2A has been a focus for development of new anticancer therapeutics, but most currently available therapeutics impact both isoforms [4].

TOP2A is a nuclear enzyme that catalyzes temporary double-stranded breaks followed by strand passage to untangle and unknot DNA [3,4]. While cells can survive without TOP2B, TOP2A appears to be essential due to the role it serves in unlinking chromosomes during replication [1]. Both isoforms contain the same domains including an N-terminal ATPase domain, a core metal-ion binding TOPRIM domain, an adjacent catalytic domain, a lower gate region, and an intrinsically disordered C-terminal domain [1,5]. The catalytic cycle involves a carefully orchestrated series of movements that are still being worked out in detail, but generally include: 1) binding of a helix-helix crossover to the core/cleavage ligation domain and within the N-terminal domain. 2) closing of the N-terminal domain as a “clamp” and cleavage of one of the segments of DNA. 3) strand passage of the intact DNA segment through the break in the cleaved segment. 4) closure of the cleaved DNA segment and religation of the break. 5) release of the passed segment from the “lower clamp” of the enzyme and opening of the N-terminal domain clamp to release the cleaved segment [4].

Interestingly, the C-terminal domains of TOP2A and TOP2B appear to be the main region where these two enzymes differ [5]. It has been proposed that the selective targeting of TOP2A may be possible through the CTD, but it will require detailed studies to understand what role or roles the CTD is playing and what regions may be suitable for targeting [4-6]. Several studies have identified key regions and functions within the CTD already including the nuclear localization signals (NLSs), chromatin tether domain (ChT), and many sites of post-translational modifications [5,7-13].

Previously, we set out to examine the CTD of TOP2A through a series of mutants of positions within the CTD. Through site-directed mutagenesis, we examined the effects of replacing Ser and Thr resides in the CTD in groups of 1-7 residues at a time [6]. While some of these mutations had little impact, some mutants were found to have decreased catalytic activity while others appeared to have increased activity. Our work demonstrated that the CTD influences catalytic activity of the enzyme, and we propose this supports the hypothesis that targeting of the CTD may be an effective strategy for regulating function of TOP2A. Combined with new bioinformatics evidence, we propose that there is significant interaction between the CTD and the N-terminal ATPase domain, which may help clarify the impact of the CTD on catalytic activity [14].

In this present brief study, we revisit several of the mutants that displayed the largest perturbations of activity to explore the impact of the CTD mutations on functions involving the N-terminal domain (Table 1). We first examine whether these enzymes have any defects in the ability to form an N-terminal clamp in the presence of ATP or AMP-PNP. Then, we examine the enzymes in the presence of a Cu(II) thiosemicarbazone, which is a known inhibitor of ATPase activity of TOP2A. Based upon our results, mutations to the CTD may impact clamp closing and the coordination of DNA cleavage. While the exact mechanism remains to be elucidated, these results continue to support the role of the CTD as a regulator of TOP2A function.

**Table 1.** CTD Mutants of TOP2A Used in this Study.

| Enzyme | Positions Mutated | Notes<sup>1</sup> |
| --- | --- | --- |
| Mutant 1 | T1272A, T1274A, T1278A, T1279A | Slightly faster relaxation, higher etoposide-induced DNA cleavage |
| Mutant 4 | S1351A, S1354G, T1358A, S1361A, S1365G | Slower relaxation; high etoposide-induced DNA cleavage |
| Mutant 7 | S1423G, S1425A, S1426G, T1427I, S1428G, T1429I, T1430A | Slightly slower relaxation |
| Mutant 9 | S1469A, T1470I, S1471A, S1474A, S1476A | Slower relaxation |
| Mutant 10 | T1487A, S1488I, S1491A,<br>S1495I | Faster relaxation |
| Mutant 11 | S1525A | Slightly faster relaxation, slightly<br>higher DNA cleavage |

<sup>1</sup> Results are from the previous study compared to WT TOP2A [6].

## 2. Results

### 2.1 N-Terminal Clamp Closing is Impacted by CTD Mutations

To investigate whether mutations in the CTD influence the N-Terminal Clamp of TOP2A, assays were performed to measure the ability of the N-terminal clamp to form a stabilized closed form, similar to previous studies [15,16]. In this assay, TOP2A is incubated with plasmid DNA along with either ATP or AMPPNP (a nonhydrolyzable ATP analog). After incubation, reaction mixtures are filtered through a nitrocellulose filter, which will strongly bind protein (TOP2A) but not DNA. If the enzyme is “clamped” around DNA at the N-terminus, the DNA will be retained in the filter. Successive washes of increasing stringency will release DNA that is weakly bound while SDS and proteinase K are used to denature and digest the protein, respectively. DNA released by the SDS and proteinase K steps is considered to be a more “stabilized” clamp. Most DNA elutes in initial low salt wash, and this data is omitted from the bar graphs for clarity.

ATP alone was used as a control, while AMP-PNP was also evaluated for an impact on clamp-closing. As seen in figure 1A, ATP induces increased clamping in mutants 1, 7 and 10 more than in WT, which is typical due to the ability of the enzyme to undergo catalysis with ATP [16]. Further, the mutants with reduced catalytic activity in our previous study (Muts 4 and 9) displayed similar clamping activity to WT with ATP.

**Figure 1.**
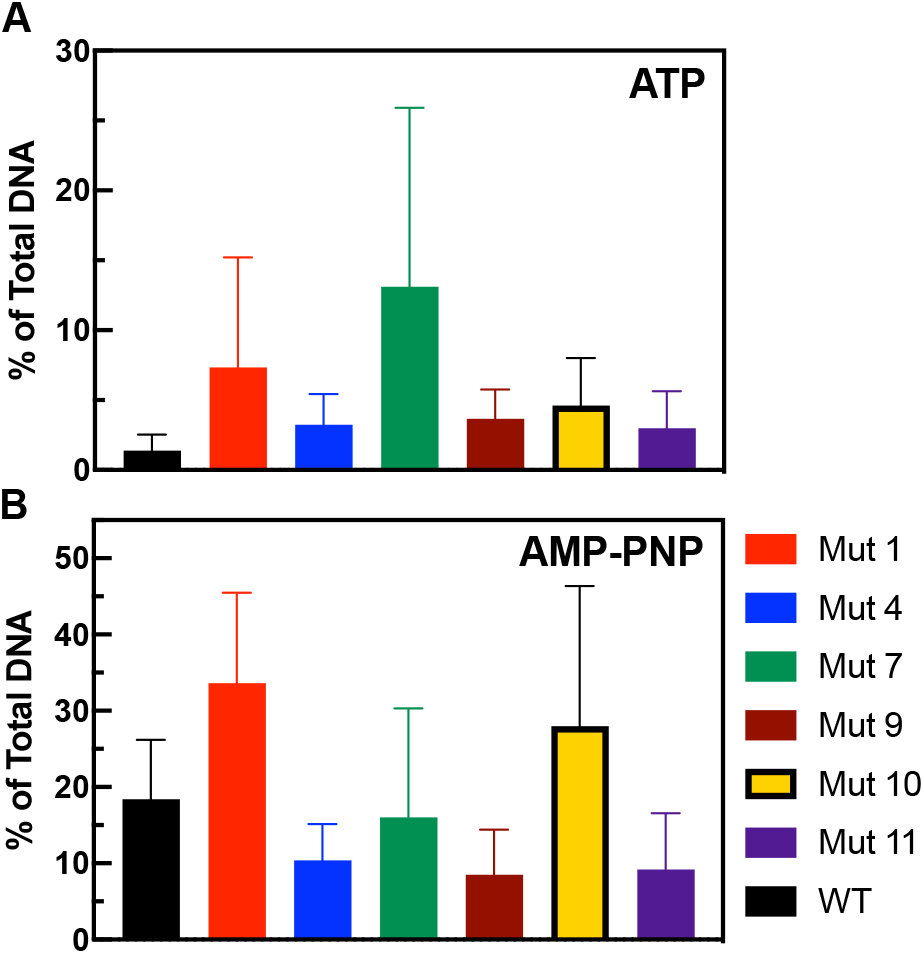
Impact of ATP and AMP-PNP on Clamp Closing with TOP2A CTD Mutants. WT and mutant TOP2A was incubated with DNA and either ATP (A) or AMP-PNP (B). DNA flow through at the SDS and Proteinase K (Pro K) wash steps are shown as a percentage of total DNA eluted. Error bars represent the standard deviation of the mean of four or more independent experiments.

In Figure 1B, AMP-PNP was used as an ATP analog that does not undergo hydrolysis, thereby stopping the enzyme in its clamped state [16]. With AMP-PNP, a larger proportion of the WT enzyme is clamped around DNA indicating that the analog has trapped the enzyme in a clamped state. Similar increases are seen with mutants 1, 7, and 10. Interestingly, it appears that mutants 4 and 9, which have the weakest catalytic activity also have decreased clamp closing with AMP-PNP. These data suggest that there may be a disruption in the catalytic function as a result of the CTD mutations.

### 2.2 Thiosemicarbazones Increase DNA Cleavage by C-Terminal Domain Mutants of TOP2A

In order to explore whether the CTD mutations impacted the mechanism of Cu-APY-ETSC, plasmid DNA cleavage assays were performed. In these experiments, increasing concentrations of Cu-APY-ETSC were incubated with plasmid DNA in the presence of WT or CTD mutants of TOP2A. As seen in Figure 2, the overall trend between mutants and WT enzyme are very similar while the specific levels of DNA cleavage enhancement vary.

**Figure 2.**
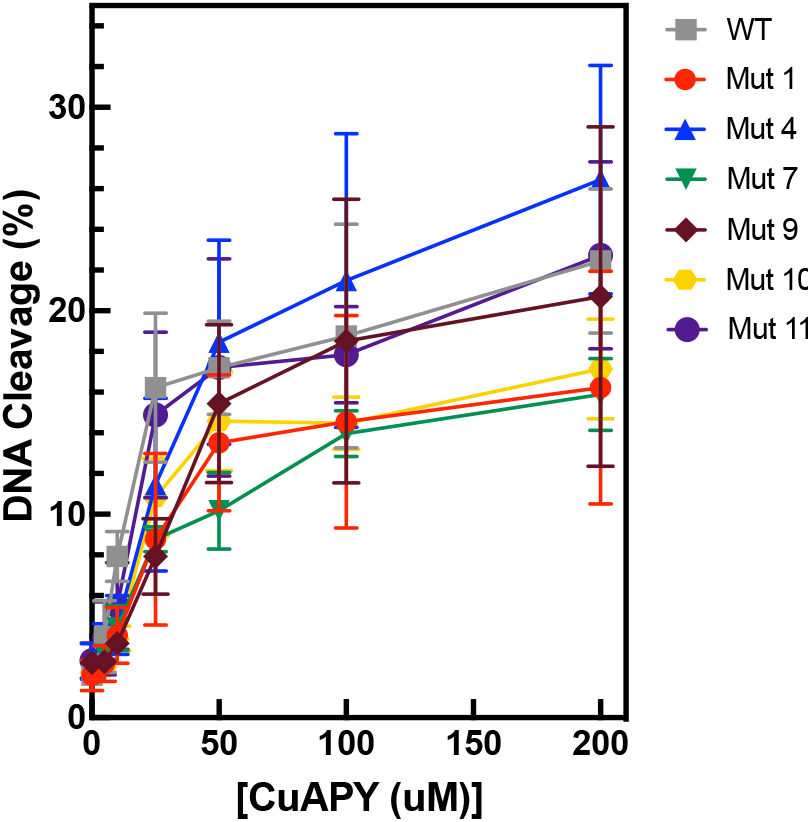
Plasmid DNA Cleavage in the Presence of Cu-APY-ETSC by WT and CTD mutant TOP2A. Reactions were incubated with increasing concentrations of Cu-APY-ETSC (CuAPY) in the presence of WT or CTD mutant TOP2A. Double-stranded DNA cleavage is quantified and plotted. Error bars represent the standard deviation of the mean of four or more independent experiments.

As seen in Figure 2, the CuAPY is able to cause increased levels of DNA cleavage with the CTD mutants similar to WT TOP2A. Despite having decreased catalytic activity in our previous study, mutants 4 and 9 display essentially WT ability to cleave DNA in the presence of CuAPY. The remaining mutants are also in range of the WT enzyme. This indicates that the impact of CuAPY is not perturbed by the CTD mutants. CuAPY is proposed based upon computational evidence to bind at or near the ATP binding pocket, this indicates that these mutations being examined may not have a major influence on this region.

### 2.3 Thiosemicarbazones Increase the DSB:SSB Ratio

Using the plasmid DNA cleavage data, a simple DSB/SSB ratio can be calculated from the level of nicked and linearized DNA in each lane. This ratio serves as a crude measurement of the level of coordination between the two active sites. Increasing levels of DSBs indicates that the DNA spends more time in a fully cleaved state. As Cu-APY-ETSC concentration increases, coordination increases in WT and all mutants (Figure 3). CTD mutants showed variability in their response to Cu-APY-ETSC. Mutants 1, 7, 10, and 11 followed the same general trend as WT though some did not quite reach the same ratio. Interestingly, mutants 4 and 9 became more coordinated as the concentrations increased rather than reaching a plateau.

**Figure 3.**
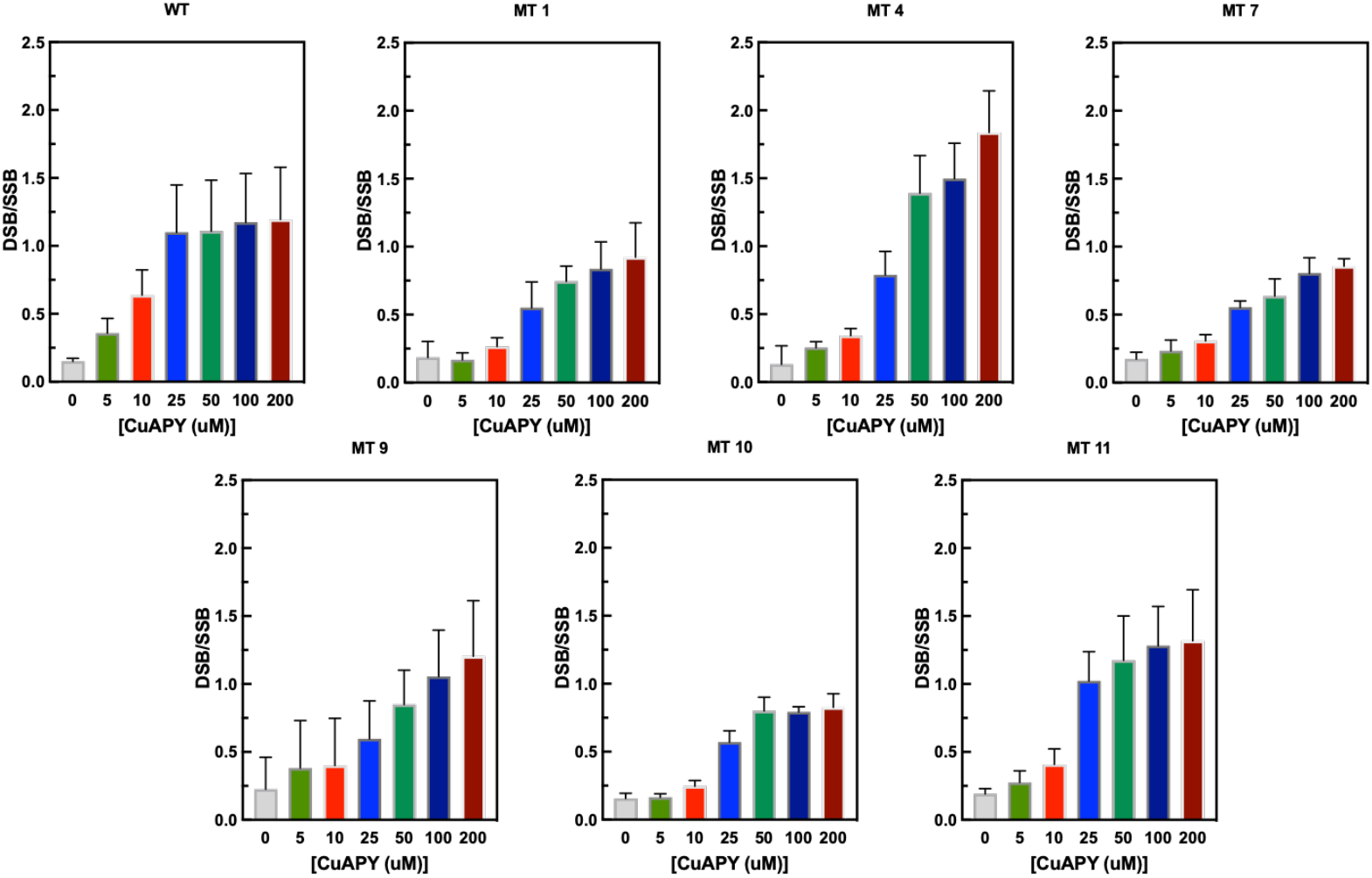
Double-stranded vs. Single-stranded Plasmid DNA Cleavage Ratios by WT and CTD Mutant TOP2A. Ratios of double-stranded to single-stranded DNA cleavage are shown for reactions with WT and CTD mutant TOP2A in the absence (0) or presence of Cu-APY-ETSC (CuAPY). Error bars represent the standard deviation of the mean of four or more independent experiments.

This is particularly noticeable in mutant 4, which reaches the highest levels of coordination.

## 3. Discussion

Increasing evidence indicates that the CTD of TOP2A serves a vital regulatory role in the function of TOP2A including interactions with other protein partners, localization on the chromatin, and even influencing catalytic activity [5,12,13,17]. Previously, we generated 11 mutants with 1-7 Ser/Thr residues in the CTD being changed to Ala/Ser/Gly [6]. We found the mutations had an impact on the activity of TOP2A in several cases, and we selected some of those mutants to follow up on in this present study.

Mutants 4 and 9 displayed decreased catalytic activity, and in the present study, these two enzymes displayed decreased clamp closing activity in the presence of AMP-PNP. Thus, it is possible that the mutated regions are having some impact on the ability of the N-terminal domain to clamp around the DNA. Mutant 4 involves mutations between amino acids 1351-1365 in the CTD, and this region includes residues that are involved in enzyme stability including phosphorylation sites and sites involved in Ub-mediated degradation [5,18-21]. Mutant 9 includes mutations between residues 1469-1476. These positions include phosphorylation sites that are modified in a cell cycle-dependent manner and are modified in association with mitosis [7,8,20,22-25]. Thus, it is possible that these regions play a role in influencing the activity of TOP2A. While both of these mutants demonstrated increases in DNA cleavage in the presence of CuAPY, they also displayed a different trend in the DSB/SSB rations: stairstep increases instead of a plateau. This effect was more pronounced in mutant 4, and implies there is an increased level of coordination between the two active sites in this mutant. The mechanism for this increase in coordination is unknown.

Mutant 1 displays higher levels of clamp closing with both ATP and AMP-PNP. Consistent with this, mutant 1 was shown previously to relax and decatenate faster than the WT enzyme. This may indicate that mutant 1 more readily associates with DNA in a catalytically active way. The region of mutant 1 is positions 1272-1279, and this includes one of the NLS regions. While this region clearly has a role in nuclear localization, it is unclear whether other roles exist, and the phosphorylation sites in this region have not been examined for their cellular effects [7,8,26].

Mutants 7 and 10 also show increases in clamp closing with AMPPNP similar to WT. Mutant 7 also shows an increase above WT levels in the presence of ATP. This result is consistent with previous data on mutant 7 where decreased rates of relaxation were observed. It is possible that ATP is not cycling through this enzyme in a normal timeframe, and the enzyme is getting stuck to some degree. Mutant 7 includes positions 1423-1430. Interestingly, our bioinformatic analysis of TOP2A sequences using PSICalc indicates an interdependency between position 1430 and positions around the ATP binding pocket (119,120,126) [14]. This computational data may help us understand the biochemical data we have collected. If position 1430 actually interacts with the positions in the N-terminal ATPase domain, specifically around the binding pocket, this mutation may have disrupted that interaction and could be affecting catalytic activity as a result.

Mutant 10 includes mutations of Ser/Thr between residues 1487 and 1495. Similar to mutant 7, residues in this range display an interdependency with residues in the ATPase domain [14]. Position S1488 clusters with residues 388-390 and S1491 clusters with residues 194, 197, and 211. While positions 388-390 are part of the transducer domain that connects the ATPase to the TOPRIM domain in the core, positions 194-211 are part of the ATPase domain. Previous data with mutant 10 indicates that this enzyme actually has increased rate of relaxation, which may reflect the impact of these interdependencies being disrupted via mutation [6]. This data cannot demonstrate whether it is the change at 1488, 1491, or both that is significant. Additional studies will be needed to clarify whether these positions are playing distinct roles in this interaction.

It should be noted that this data cannot clarify whether the mutation impacts a direct interaction between the mutated region of the CTD and the ATPase or if the disruption caused by the mutations is impacted another CTD region that interacts with the ATPase. However, the combination of biochemical and computational evidence support the concept that the CTD and ATPase interact in some way.

## 4. Materials and Methods

### 4.1 Enzymes

Wild-type TOP2A and mutant TOP2A expression was carried out in Saccharomyces cerevisiae JEL1Δtop1 using pESC-URA-TOP2A as previously described [6]. CTD mutants were generated by Genscript as described in Dougherty, et. Al [6]. Protein samples were stored as a 1 or 1.5 mg/mL (3−4 μM) stock in 50 mM Tris-HCl, pH 7.7, 0.1 mM EDTA, 750 mM KCl, and 5% glycerol at −80 °C.

### 4.2 Plasmid DNA Preparation

Negatively supercoiled pBR322 DNA was purified from DH5α utilizing a Plasmid Mega Kit (Qiagen, Hilden, Germany). Negatively supercoiled pHOT-1 DNA was purified from DH5α utilizing a Qiagen Plasmid Midi Kit (Qiagen, Hilden, Germany).

### 4.3 Compounds

AMP-PNP was obtained from Roche Diagnostics (Mannheim, Germany) and stored at a concentration of 20 mM in water at −20°C. ATP was obtained from Sigma (St. Louis, MO) and stored at a concentration of 20 mM in water at −20°C. The compound N4-ethylthiosemicarbazone (APY-ETSC) was synthesized from its parent thiosemicarbazide as described previously [27]. The copper (II) complex ([Cu-(APY-ETSC)Cl]) was prepared by heating the TSC with CuCl_2_ [27,28]. The Cu-APY-ETSC was stored at 4°C in 100% DMSO at a concentration of 20mM [29].

### 4.4 N-Terminal Clamp Closing Assay

Nitrocellulose filters were placed into filter baskets and incubated at room temperature in 100 mM KCl with salmon testis DNA (Sigma, St. Louis, MO) and acetylated BSA (Promega, Fitchburg, WI) for 20 minutes before low speed centrifugation and an additional wash with a buffer solution (50 mM Tris-HCl, pH 8.0, 100 mM KCl, 1 mM EDTA, and 8 mM MgCl_2_). Reaction mixtures contained ∼ 80 nM TOP2A, ∼8 nM pHOT1 plasmid DNA, in the reaction buffer solution mentioned above. The reaction mix was then incubated for 5 minutes at 37°C. Next, 2 mM ATP or AMP-PNP was added to the master mix and was incubated for an additional 5 minutes at 37°C. The reaction mix was then loaded into the filter baskets with nitrocellulose filters and centrifuged at 1,000 rpm for 60s. Filters were washed with the reaction buffer mentioned above (“Low salt”), then a 1 M NaCL solution (“High Salt”), and finally a SDS wash (10 mM Tris-HCl, pH 8.0, 1 mM EDTA, and 0.5% SDS) heated to 65 °C. Flow through was collected at each wash. Filters were soaked in a Proteinase K (Amresco, Solon, OH) solution for 10 min at 45°C before a final spin into a clean tube. Low Salt, High Salt, and SDS samples were concentrated using the Monarch PCR & DNA Cleanup Kit (NEB, South Hamilton, MA). Samples were resolved on a 1% agarose TAE gel with ethidium bromide and imaged using BioRad ChemiDoc MP Imaging system (Hercules, CA). Data was plotted and analyzed using GraphPad Prism 6 (La Jolla, CA).

### 4.5 Plasmid DNA Cleavage Assay

Plasmid DNA Cleavage assays were performed using conditions described previously [6,30]. Reactions were performed in the presence of 0-200 μM concentration of CuAPY. WT or mutant TOP2A and 5 nM pBR322 DNA in a buffer of 20 μL of 10 mM Tris-HCl, pH 7.9, 100 mM KCl, 1 mM EDTA, 5 mM MgCl_2_, and 2.5% glycerol were incubated for 6 minutes at 37°C. EcoR1 was used to form a linearized DNA control. Reactions were stopped using 2 μL of 2.5% SDS. After this, 2 μL of Na_2_EDTA at 250mM and Proteinase K at 0.8 mg/mL were added. Samples were then incubated at 45 °C for 30 minutes to degrade protein. After this, gel electrophoresis was performed on samples utilizing a 1% TAE gel with ethidium bromide, and bands were visualized using ChemiDoc MP Imaging System and Image Lab software (Bio-Rad, Hercules, CA). Graph Pad Prism 9 (San Diego, CA) was used to quantify data. Quantified data for linearized, nicked, and supercoiled bands were logged. DSB/SSB ratio was determined by dividing the linearized band by the nicked band.

## Supplementary Materials

NA

## Author Contributions

Conceptualization, J.E.D.; methodology, J.E.D.; investigation, J.R.M., L.A.F., C.T.D., D.C.E, E.B.; resources, J.E.D., X.J., E.C.L.; data curation, J.E.D., L.A.F, J.R.M; writing—original draft preparation, J.E.D, D.C.E., L.A.F, J.R.M.; writing—review and editing, J.E.D., J.R.M., L.A.F., D.C.E., C.T.D.; visualization, J.E.D., L.A.F., D.C.E., J.R.M.; supervision, J.E.D.; project administration, J.E.D.; funding acquisition, J.E.D. All authors have read and agreed to the published version of the manuscript.

## Institutional Review Board Statement

Not applicable.

## Informed Consent Statement

Not applicable.

## Data Availability Statement

Data is available upon reasonable request to the corresponding author.

